# Reference-free resolution of long-read metagenomic data

**DOI:** 10.1101/811760

**Authors:** Lusine Khachatryan, Seyed Yahya Anvar, Rolf H. A. M. Vossen, Jeroen F. J. Laros

**Affiliations:** Department of Human Genetics, Leiden University Medical Center, Leiden, The Netherlands; Leiden Genome Technology Center, Leiden University Medical Center, Leiden, The Netherlands; Department of Clinical Genetics, Leiden University Medical Center, Leiden, The Netherlands; GenomeScan, Leiden, The Netherlands

**Keywords:** Metagenomics binning, PacBio sequencing, metagenome resolving

## Abstract

**Background:** Read binning is a key step in proper and accurate analysis of metagenomics data. Typically, this is performed by comparing metagenomics reads to known microbial sequences. However, microbial communities usually contain mixtures of hundreds to thousands of unknown bacteria. This restricts the accuracy and completeness of alignment-based approaches. The possibility of reference-free deconvolution of environmental sequencing data could benefit the field of metagenomics, contributing to the estimation of metagenome complexity, improving the metagenome assembly, and enabling the investigation of new bacterial species that are not visible using standard laboratory or alignment-based bioinformatics techniques.

**Results:** Here, we apply an alignment-free method that leverages on *k*-mer frequencies to classify reads within a single long read metagenomic dataset. In addition to a series of simulated metagenomic datasets, we generated sequencing data from a bioreactor microbiome using the PacBio RSII single-molecule real-time sequencing platform. We show that distances obtained after the comparison of *k*-mer profiles can reveal relationships between reads within a single metagenome, leading to a clustering per species.

**Conclusions:** In this study, we demonstrated the possibility to detect substructures within a single metagenome operating only with the information derived from the sequencing reads. The obtained results are highly important as they establish a principle that might potentially expand the toolkit for the detection and investigation of previously unknow microorganisms.

## INTRODUCTION

The analysis of metagenomics data is becoming a routine for many different research fields, since it serves scientific purposes as well as improves our life quality. Particularly, with the use of metagenomics a large step was made towards the understanding of the human microbiome and uncovering its real composition and diversity [1–6]. The understanding of the human microbiome in health and disease contributed to the development of diagnostics and treatment strategies based on metagenomic knowledge [7–14]. The study of microbial ecosystems allows us to predict the possible processes, changes and sustainability of particular environments [15, 16]. Genes isolated from uncultivable inhabitants of soil metagenomes are being successfully utilized, for example, in the biofuel industry for production and tolerance to byproducts [17–19]. Various newly discovered biosynthetic capacities of microbial communities benefit the production of industrial, food, and health products, as well as contribute into the field of bioremediation [20–23].

Despite all the progress made in resolving genetic data derived from environmental samples, it is still a challenging task. Reads binning is one of the most critical steps in the analysis of metagenomics data. To estimate the composition of a particular microbiome, it is important to ensure that sequencing reads derived from the same organism are grouped together. Currently, alignment of DNA extracted from an environmental sample to a set of known sequences remains the main strategy for metagenomics binning [24, 25]. There is a full range of techniques allowing the comparison of metagenomic reads to a reference database. It can be performed using different metagenomic data types (16S or WGS) and various matching approaches (classic alignment or use of *k*-mers or taxonomical signatures). Most of the time, the binning is performed for all reads in the database, but in some cases only a particular subset of sequencing data is selected for binning. Lastly, there is a wide spectrum of databases that can be used to perform the binning. The database might contain all possible annotated nucleotide/protein sequences, marker genes for distinct phylogenetic clades, sequencing signatures specific to particular taxa, etc. The obvious downside of all listed strategies is the incapability to perform an accurate binning for the reads of organisms that are not present in the reference database.

Metagenomic binning was improved by alignment-free approaches, which can be split into two subgroups: reference-dependent and reference-independent methods. The tools from the first subgroup utilize existing databases to train a supervised classifier for the reads binning. Various techniques can be performed to achieve this goal: linear regression, Interpolated Markov Models, Gaussian Mixture Models, Hidden Markov Models [26–32]. Even though these approaches are reference dependent, they can be used to classify reads that are derived from previously unknown species. However, the accuracy of reference-dependent methods will be always limited by the content of reference databases. The content of the current reference databases utilized for training differs from the true distribution of microbial species on our planet [33–39]. For some metagenomic datasets the amount of unknown sequences might be quite high [40, 41], thus using supervised classification tools based on known genetic sequences is questionable in such cases.

Reference-independent approaches for metagenomics binning try to solve the problem of missing taxonomic content: they are designed to classify reads into genetically homogeneous groups without utilizing any information from known genomes. Instead, they use only the features of the sequencing data (usually *k*-mer distributions, DNA segments of length *k*) for classification. One of those tools, LicklyBin, performs a Markov Chain Monte Carlo approach based on the assumption that the *k*-mer frequency distribution is homogeneous within a bacterial genome [42]. This tool performs well for very simple metagenomes with significant phylogenetic diversity within the metagenome, but it cannot handle genomes with more complicated structure such as those resulting from horizontal gene transfer [43]. Another one, AbundanceBin [44], works under the assumption that the abundances of species in metagenome reads are following a Poisson distribution, and thus struggles analyzing datasets where some species have similar abundance ratios. MetaCluster [45] and BiMeta [46] address this problem of non-Poisson species distribution. However, for these tools it is necessary to provide an estimation of the final number of clusters, which cannot be done for many metagenomes without any prior knowledge. Also, both MetaCluster and BiMeta are using a Euclidian metric to compute the dissimilarity between *k*-mer profiles, which was shown to be influenced by stochastic noise in analyzed sequences [47]. Another recent tool, MetaProb, implements a more advanced similarity measure technique and can automatically estimate the number of read clusters [48]. This tool classifies metagenomic datasets in two steps: first, reads are grouped based on the extent of their overlap. After that, a set of representing reads is chosen for each group. Based on the comparison of the *k*-mer distributions for those sets, groups are merged together into final clusters. Even though MetaProb outperformed other tools during the analysis of simulated data, it was shown to perform not very well on the real metagenomics data.

In this article we present a new technique for alignment- and reference-free classification of metagenomics data. Our approach is based on a pairwise comparison of *k*-mer profiles calculated for each sequencing read in a long-read metagenomics dataset, using the previously described kPAL toolkit [49]. It also performs unsupervised clustering to facilitate the identification of genetically homogeneous groups of reads present in a sample. The main assumption of our method is that after assigning the pairwise distances for all reads in the dataset, those belonging to the same organism will form dense groups, and thus the metagenome binning could be resolved using density-based clustering. We developed an algorithm which automatically detects the regions with high density and hierarchically splits the dataset until there is one dense region per cluster. The approach is designed to work with long reads (more than 1000 bp) since we calculate *k*-mer profiles for each read separately and shorter reads would yield non-informative profiles. We performed our analysis on long PacBio reads that were either simulated or generated from a real metagenomic sample. We have shown that despite the fact that PacBio data is known to have a high error rate, the approach successfully performed read classification for simulated and real metagenomic data.

## MATERIALS AND METHODS

### 1. Software

All analyses were done using publicly available tools (parameters used are listed below for each specific case) along with custom Python scripts.

### 2. PacBio data simulation

Complete genomes of five common skin bacteria were used to generate artificial PacBio metagenomes (Table 1). The reads were simulated from reference sequences using the PBSIM toolkit [50] with CLR as the output data type and a final sequencing depth of 20. For the calibration of the read length distribution, a set of previously sequenced *C. difficille* reads [51] was used as a model.

**Table 1.** Content of artificial metagenomics PacBio datasets.

| Reads origin | RefSeq AC | Genome length, Mb | Number of reads per dataset |  |  |  |
| --- | --- | --- | --- | --- | --- | --- |
|  |  |  | 0% noise | 5% noise | 10% noise | 15% noise |
| *S. mitis* | NC_013853.1 | 2.1 | 1 246 | 1 183 | 1 121 | 1 059 |
| *P. acnes* | NC_017550.1 | 2.5 | 1 443 | 1 371 | 1 298 | 1 226 |
| *S. epidermidis* | NC_004461.1 | 2.6 | 1 448 | 1 376 | 1 304 | 1 231 |
| *A. calcoaceticus* | NC_016603.1 | 3.9 | 2 236 | 2 125 | 2 013 | 1 901 |
| *P. aeruginosa* | NC_002516.2 | 6.3 | 3 627 | 3 446 | 3 264 | 3 083 |

### 3. Bioreactor metagenome PacBio sequencing

Bioreactor metagenome coupling anaerobic ammonium oxidation (Annamox) to Nitrite/Nitrate dependent Anaerobic Methane Oxidation (N-DAMO) processes [52] was used to generate WGS PacBio sequencing data.

Metagenome contained the N-DAMO bacteria *Methylomirabilis oxyfera* (complete genome with GeneBank Acsession FP565575.1 was used as a reference), two Annamox bacteria (*Kuenenia stuttgartiensis*, assembly contigs from the Bio Project PR-JEB22746 were used as a reference and a member of *Broccardia* genus, assembly contigs of *Broccardia sinica* from Bio Project PRJDB103 were used as reference) and an archaea species *Methanoperedens nitroreducens* (assembly contigs from the Bio Project PRJNA242803 were used as a reference).

Bacterial cell pellets were disrupted with a Dounce homogenizer. DNA was isolated using a Genomic Tip 500/G kit (Qiagen) and needle sheared with a 26G blunt end needle (SAI Infusion). Pulsed-field Gel electrophoresis was performed to assess the size distribution of the sheared DNA. A SMRTbell library was constructed using 5µg of DNA following the 20kb template preparation protocol (Pacific Biosciences). The SMRTbell library was size selected using the BluePippin system (SAGE Science) with a 10kb lower cut-off setting. The final library was sequenced with the P6-C4 chemistry with a movie time of 360 minutes.

### 4. Reads origin checking

Reads were corrected using the PacBio Hierarchical Genome Assembly Process algorithm before being mapped to the genomes of the references of expected metagenome inhabitants using the BLASR aligner [53] with default settings. The alignments were used to determine the origin of the reads. Reads that were not mapped during the previous step were subjected to the BLASTn [54] search against the NCBI database. The identity cut-off was set to 90, the (E)value was chosen to be 0.001.

### 5. Bioreactor metagenome PacBio reads assembly

The assembly of corrected PacBio reads was performed using the FALCON [55] assembler. The resulting contigs were mapped to the candidate reference genomes using LAST [56] with default settings. To determine the similarity cutoff for the mapping procedure, the curve representing the number of contigs versus the similarity to the reference genome was analyzed. The first inflection point at (in case of mapping contigs to the *M.oxyfera* genome 12%), dividing the fast-declining part of the curve from the slow-declining part, was chosen as a threshold (See Section S1 of Additional file 1 for more details).

### 6. Binning procedure

For each read, the frequencies of all possible five-mers are calculated using the *count* command of the kPAL toolkit. The resulting profiles are balanced (a procedure that compensates for differences that occur because of reading either the forward or reverse complement strand) and compared in a pairwise manner by using the *balance* and *matrix* commands of kPAL accordingly, yielding a pairwise distance matrix. Normalization for differences in read length is dealt with by the scaling option during the pairwise comparison.

The resulting distance matrix, hereafter called the original distance matrix, was subjected to a multi-step clustering procedure. A schematic representation of this procedure can be found in Fig. 1. Due to practical limitations (runtime), this analysis was restricted to a set of 10 000 randomly selected reads.

This multi-step clustering procedure works recursively: it starts with the analysis of a set of reads and either reports the entire set as one cluster, or it splits the set into two subsets, which are each analyzed using the same procedure. The decision whether to split the set of reads into two subsets is made using the following approach. First, the pairwise distances for all reads in the set are extracted from the original distance matrix in order to construct the working distance matrix. After that, the dimensionality of the analyzed set is decreased to three using the t-SNE algorithm [57] in order to reduce noise caused by outliers in the distance matrix. The reads, now represented by a point in three-dimensional space, are subjected to density-based clustering using the DBSCAN algorithm [58] with the default distance function. We choose the *MinPts* parameter of DBSCAN (the minimal amounts of points in the neighborhood to extend the cluster) to be either 1% of the size of the dataset for sets larger than 2000 reads, or 20 for sets smaller than 2000 reads. The number of clusters found by DBSCAN depends on the neighborhood diameter *ε*. When *ε* is too small, no clusters are reported since all points are isolated. On the other hand, when *ε* is too large all points are grouped into one cluster. Our algorithm therefore performs a parameter sweep for *ε*, from the value providing zero clusters to the value with which 99% of the reads are grouped in one cluster for the chosen *MinPts*.

**Fig. 1.**
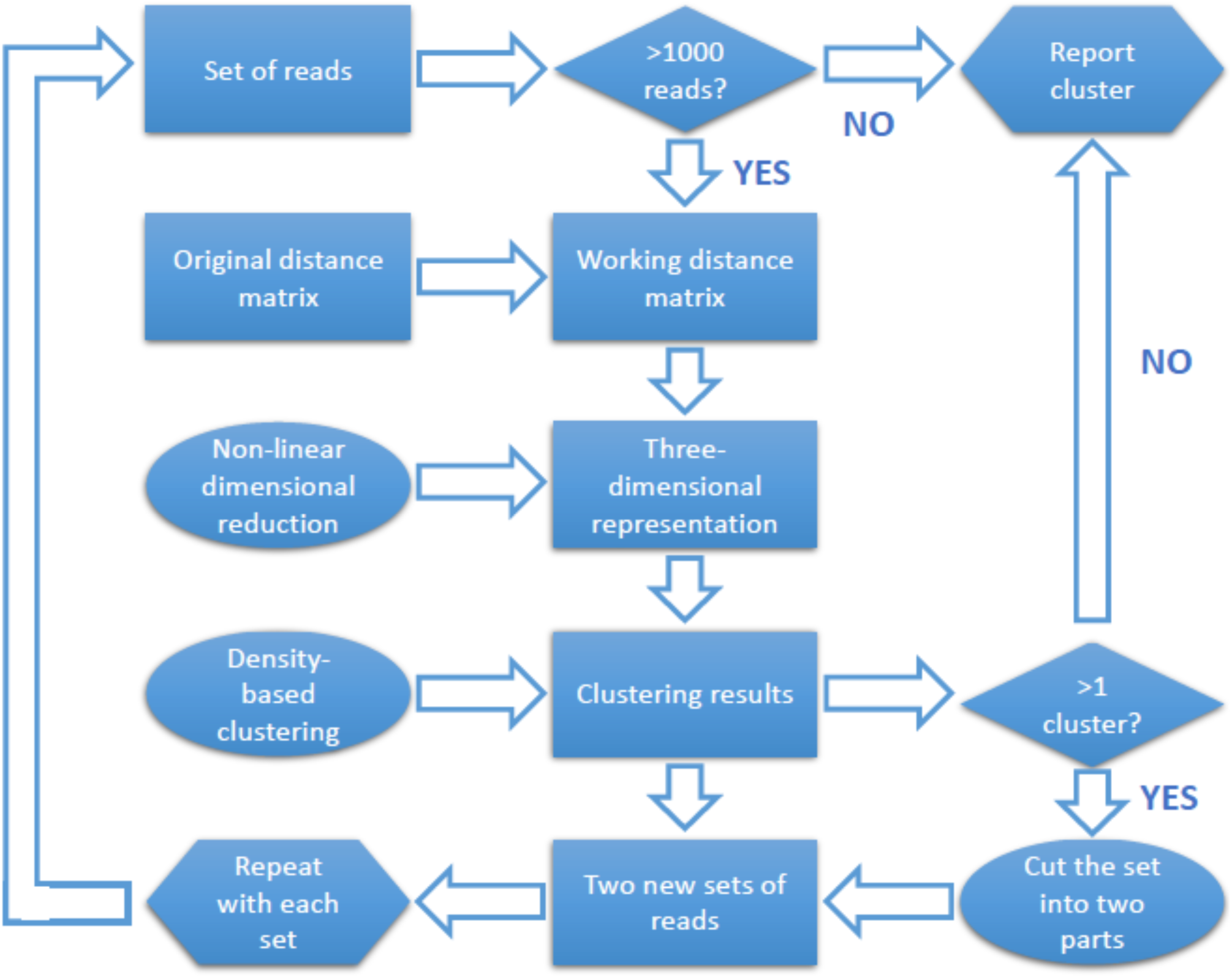
Schematic representation of the clustering procedure.

The results of this parameter sweep are used to check the dependency of the number of dense clusters on a particular *ε* (only clusters larger than 100 points are considered) and how many points of the analyzed set are included in the obtained clusters (Fig. 2). If for some *ε* there are two or more clusters that together cover more than half of the total amount, the analyzed set is divided into two new sets (Fig. 2A). The analyzed set is reported as one cluster if the aforementioned condition is not satisfied (Fig. 2B), or when the size of the analyzed set was smaller than 1000 points.

**Fig. 2.**
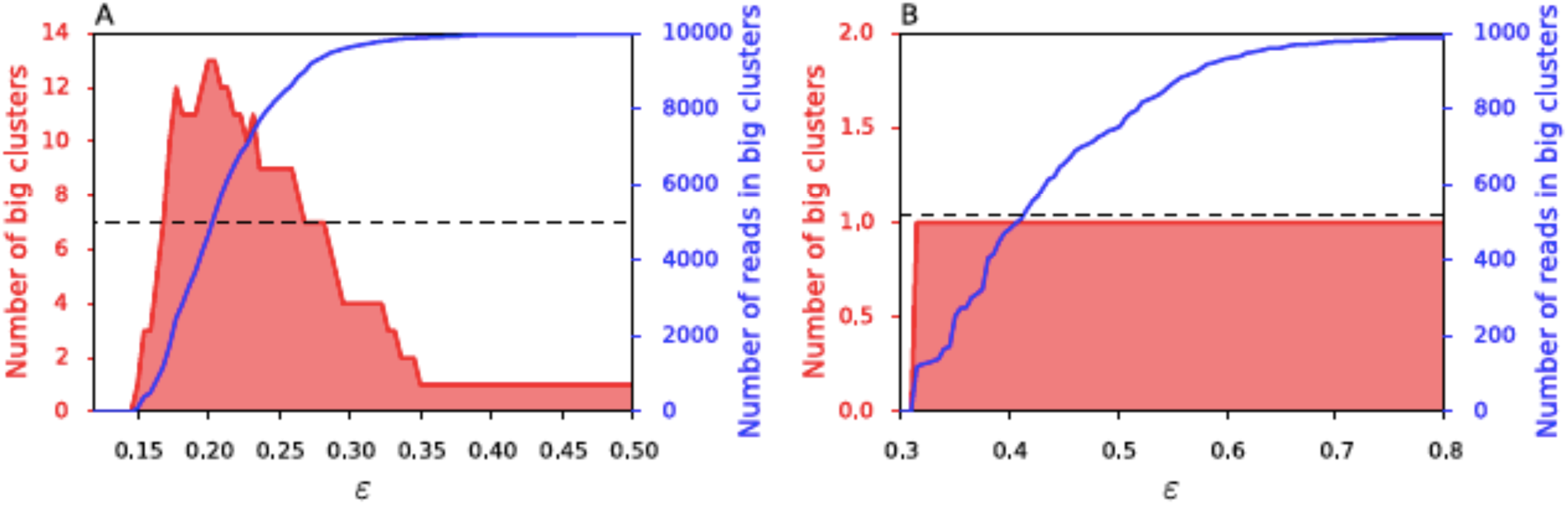
Density-based clustering analysis example. The data is clustered with DBSCAN with ε ranging from 0 to the value when 90% of the points are assigned to one cluster. When at least half of the data set is assigned to a dense cluster, the number of clusters is used to determine whether subdivision of the data set is required. Only if more than one cluster is identified at this point, the procedure is repeated recursively with two partitions of the data. The partitions are determined by using the largest ε that clusters the data into two clusters. In this example two datasets are shown: one that was further split into two partitions (A) and one that was reported as one dense cluster (B).

The division is done using the following strategy. DBSCAN is performed using the optimal *ε*, yielding two dense clusters that serve as center points for two partitions. Each of the remaining unclassified points is assigned to the cluster containing the closest classified neighbor.

### 7. Classification for larger sets

Read classification for sets larger than 10 000 was performed in two steps. First, 10 000 reads (larger than 10kb) were randomly chosen and classified using the algorithm described in previous section. After that, the pairwise distances between every unclassified read and every classified read were calculated using their 5-mer profiles. These distances were used to assign the unclassified read to the cluster containing the closest classified read.

### 8. Data availability

Sequencing reads of bioreactor metagenome were submitted to NCBI under the BioProject number PRJNA487927. Artificial PacBio metagenomic reads with the addition of 0%, 5%, 10%, and 15% of real “noise” reads were submitted to NCBI under the BioProject number PRJNA533970. Supplementary materials were deposited on Figshare and available for downloading using the following link: https://doi.org/10.6084/m9.figshare.c.4218857.v1.

Example of the classification procedure can be found using the following link: https://git.lumc.nl/l.khachatryan/pacbio-meta/blob/master/analysis/real_data/tsne_subset2/analysis_example.ipynb

## RESULTS

### 1. Reads classification in artificial PacBio metagenomes

To construct artificial metagenomes, we used simulated PacBio reads based on the genomes of five common skin flora bacteria together with so-called “noise” reads. These are reads from a PacBio sequencing data of an environmental metagenome [59] that were not assigned to the major inhabitant *K. stuttgartiensis* or other known organisms. They were added to represent low abundant species that are present in any typical metagenomic dataset.

We constructed four artificial PacBio datasets in this way, each containing 10 000 randomly selected reads (length > 9kb) containing 0%, 5%, 10% and 15% noise reads, respectively. For the simplicity the number of simulated reads was adjusted to provide an equal abundance for each bacterium in the final metagenome (see Table 1).

We subjected each dataset to the classification procedure described in Section 6 of MATERIALS AND METHODS. The reads in the resulting clusters were then classified according to their origin (See Section S2 of Additional file 1 for more data).

In Fig. 3, it can be seen that for each experiment we obtained five large clusters (> 1 000 reads) consisting mainly of reads belonging to the same species.

**Fig. 3.**
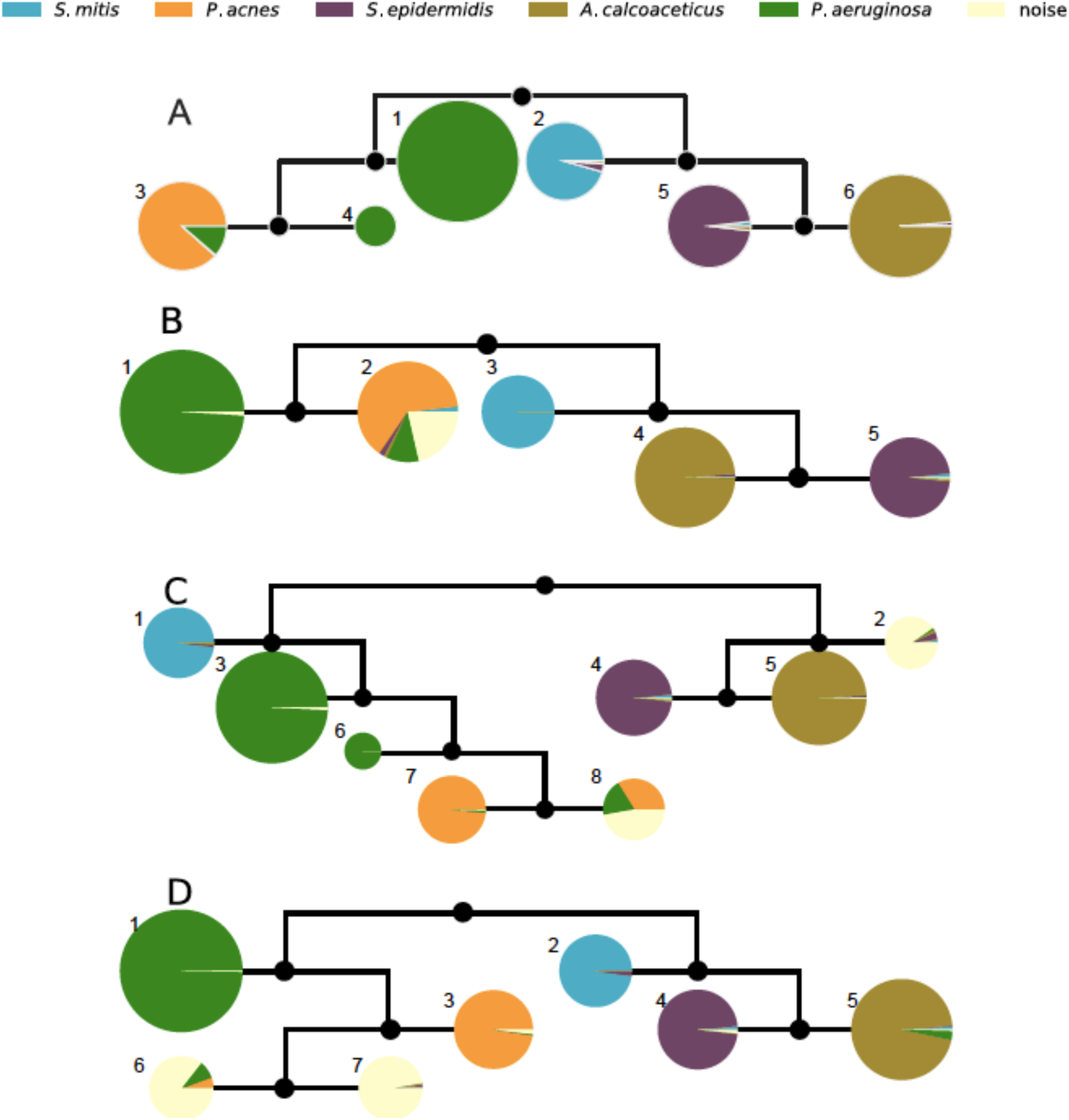
Classification recall for artificial PacBio metagenomes. Subsets that were subjected to the partitioning are shown as black circles, final clusters are represented as pie charts with the color indicating the reads origin. The area of the pie chart corresponds to the relative cluster size. The cluster number is shown next to each pie chart. The results are shown for datasets with 0% (A), 5% (B), 10% (C) and 15% (D) of noise reads.

For all three datasets containing noise reads we see the tendency of noise reads to be clustered with some fraction of *P. acnes* and *P. aeruginosa* reads.

However, as can be seen from Fig. 3 and Table 2, increasing the noise content leads to better isolation of these reads. Indeed, for dataset B (5% of the noise reads), the majority of noise reads were assigned to the cluster that is primarily occupied by reads belonging to *P. acnes and P. aeruginosa.* Increasing the noise content (dataset C and D in Fig. 4, 10% and 15% noise reads accordingly) led to the appearance of two clusters which contain mostly noise reads (Table 2, A).

**Fig. 4.**
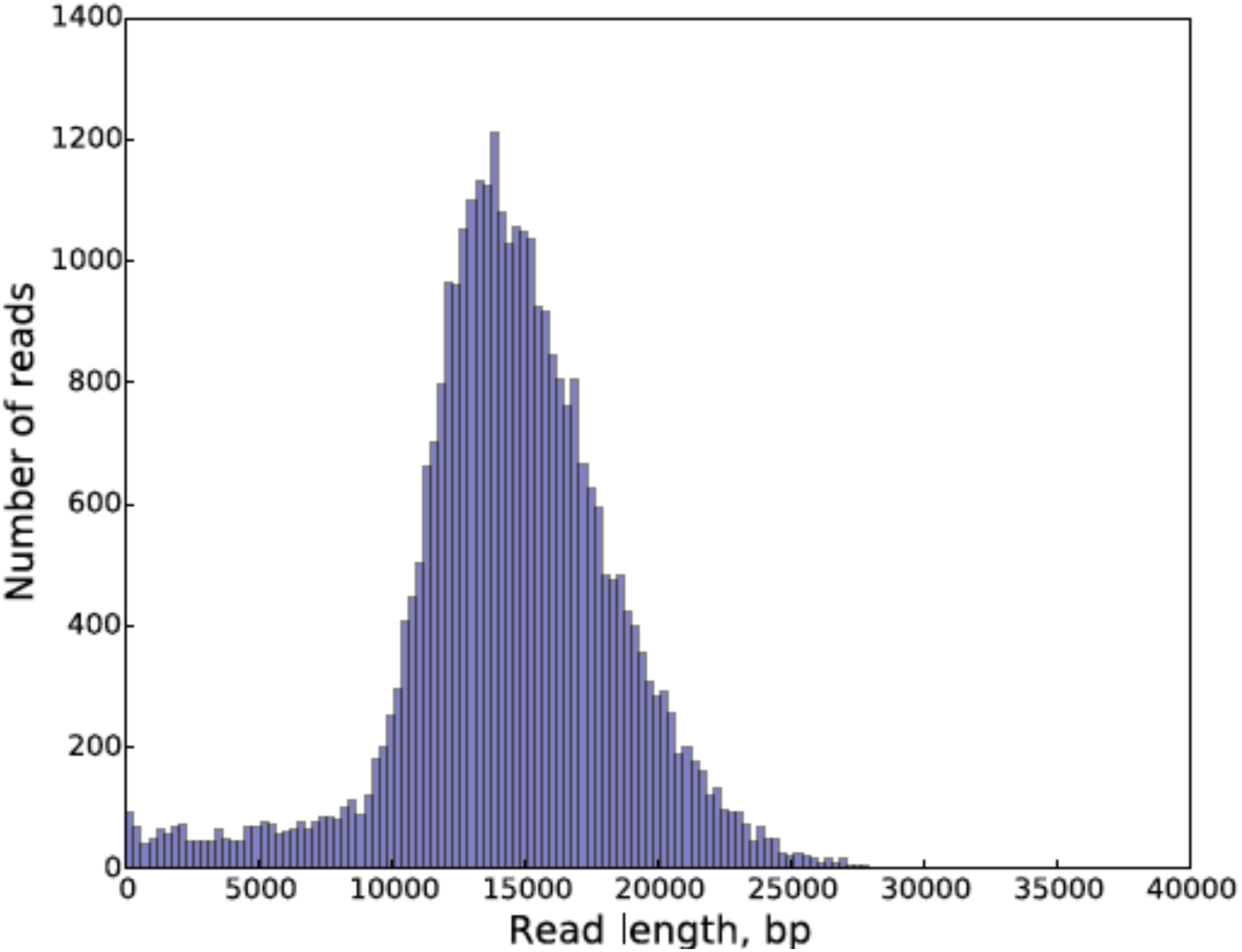
Bio reactor metagenome reads length distribution.

**Table 2.**
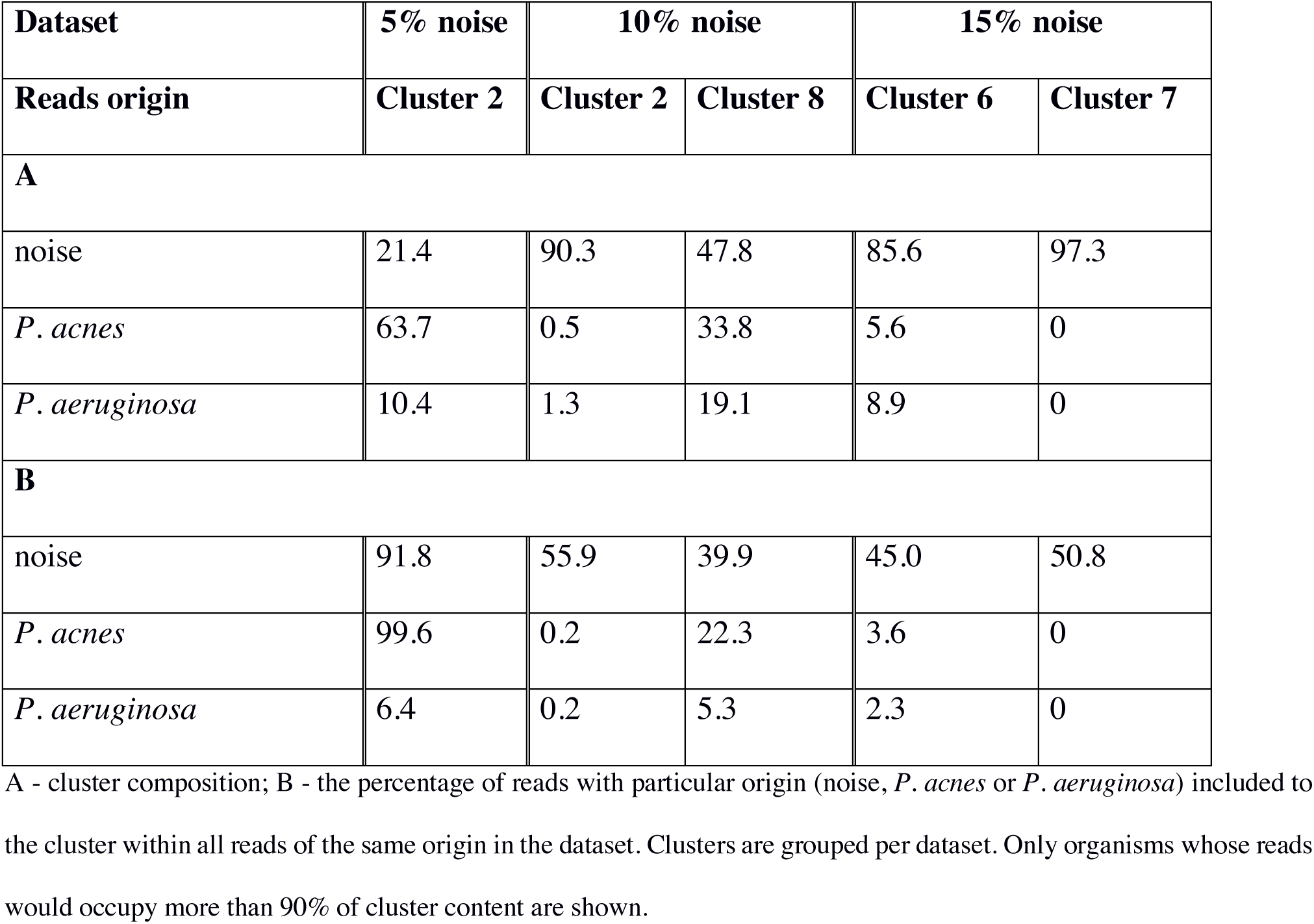
Composition of clusters containing the majority of noise reads after the classification procedure for three artificial PacBio datasets.

| Dataset | 5% noise | 10% noise |  | 15% noise |  |
| --- | --- | --- | --- | --- | --- |
| Reads origin | Cluster 2 | Cluster 2 | Cluster 8 | Cluster 6 | Cluster 7 |
| **A** |  |  |  |  |  |
| noise | 21.4 | 90.3 | 47.8 | 85.6 | 97.3 |
| *P. acnes* | 63.7 | 0.5 | 33.8 | 5.6 | 0 |
| *P. aeruginosa* | 10.4 | 1.3 | 19.1 | 8.9 | 0 |
| **B** |  |  |  |  |  |
| noise | 91.8 | 55.9 | 39.9 | 45.0 | 50.8 |
| *P. acnes* | 99.6 | 0.2 | 22.3 | 3.6 | 0 |
| *P. aeruginosa* | 6.4 | 0.2 | 5.3 | 2.3 | 0 |

A - cluster composition; B - the percentage of reads with particular origin (noise, *P. acnes* or *P. aeruginosa*) included to the cluster within all reads of the same origin in the dataset. Clusters are grouped per dataset. Only organisms whose reads would occupy more than 90% of cluster content are shown.

We also see that with the *increase* of noise content, the fractions of *P. acnes* and *P. aeruginosa* reads included in the same clusters as the noise reads are dropping (Table 2, B). In conclusion, the more noise reads were added to the dataset, the more they were grouped together in one or two clusters (Table 2, A).

### 4.2 PacBio sequencing of bio reactor metagenome

After sequencing and correction, we obtained 31,757 reads longer than 1kb for the bio reactor metagenome. The read length distribution for this dataset can be found in Fig. 4.

Reads were mapped to the genomes of the expected metagenome inhabitants or genomes of closely related species. Since the groups of reads that we could map to the genomes of *K. stuttgartiensis* and *B. sinica* had a significant overlap (27%), we decided to combine reads mapped to the reference genomes of these two organisms in one group. We detected almost no (0.01%) reads that would map to the *M. nitroreducens* genome in the sequencing data, suggesting that this organism was either not present in the metagenome sample, or that its DNA could not be isolated reliably during the sample preparation. Thus, we divided our reads into three groups: uniquely mapped on *M. oxyfera* (4,903 reads), uniquely mapped on *K. stuttgartiensis/B. sinica* (2973 reads), and all remaining reads with unknown origin (∼75%, 23881 reads). The reads with unknown origin were checked with the BLASTn software against NCBI microbial database, to find significant similarity to any known organism. However, only 334 reads (less then 2% of total number of checked reads) got hits; there were no organisms among the obtained hits reported more than 53 times.

### 4.3 Bio reactor metagenome PacBio read classification

For the reads originating from *M. oxyfera* and *K. stuttgartiensis/B. sinica*, we checked whether the data was clustered by origin. Since roughly 75% of this sequencing data is of unknown origin, we assessed whether the clustering results for reads with unknown origin is robust. To do this, we created five subsets using the bio reactor metagenome sequencing data. Each subset contains 10,000 randomly selected reads with length > 10kb. After subjecting each subset to the classification procedure, we checked whether reads, shared by two subsets, are being clustered similarly. We compared all clusters from different subsets in a pairwise manner and marked two clusters ’similar’ when they shared at least 25% of their content. On average, every pair of subsets shared 34% of their content. Thus, in case of perfect matching of clustering results, the pair of clusters from two different subsets should on average share 34% of their content. The 25% cutoff value was chosen to compensate for possible flaws introduced by clustering mis-assignments. In Fig. 5 this analysis is shown as a graph: each pie chart represents a cluster obtained for one of the subsets (with a subset number marked next to the pie chart).

**Fig. 5.**
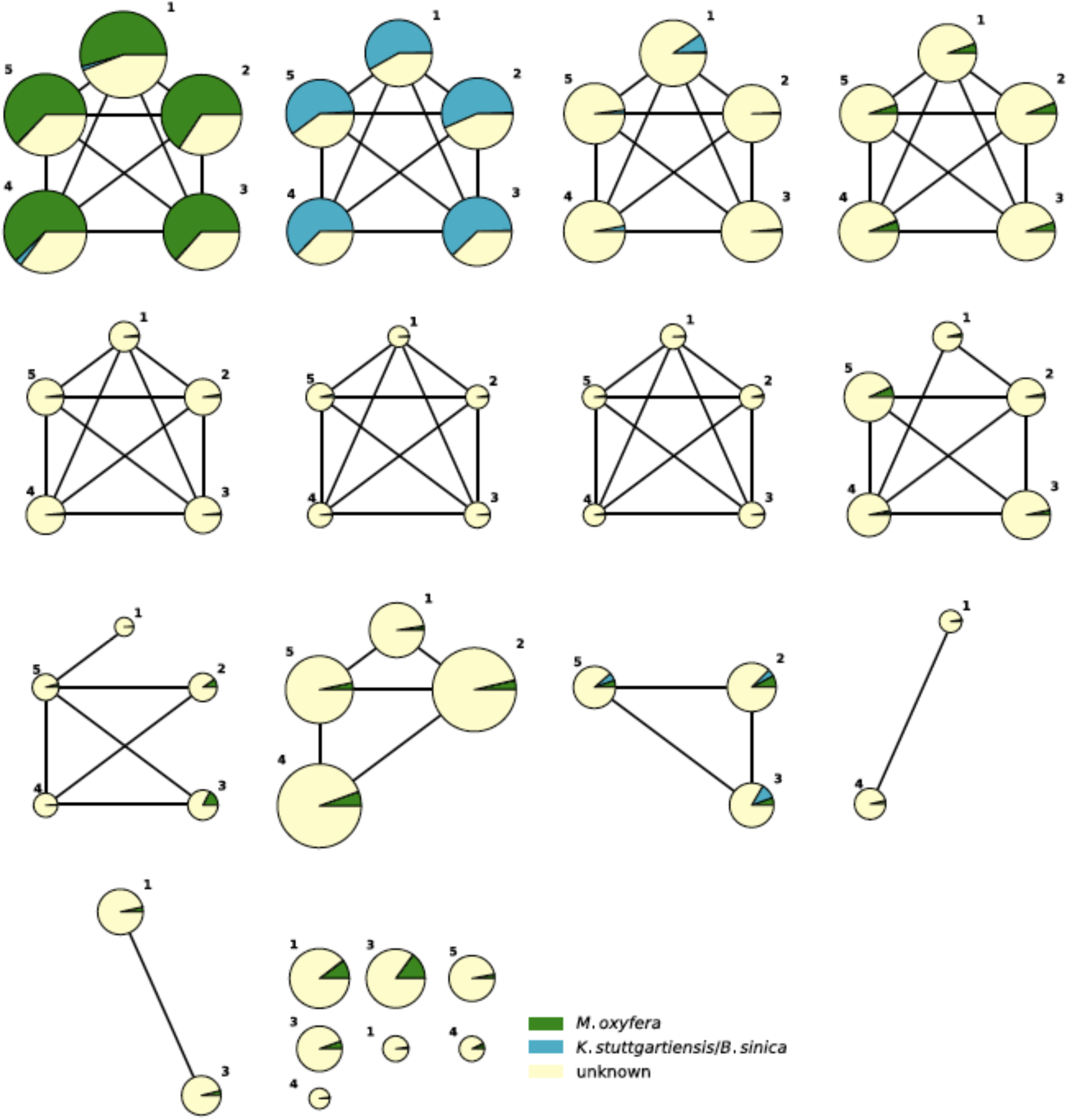
Comparison of classification results obtained for five Bio reactor sub-datasets. The pie charts represent reported clusters for all sub-datasets colored by the origin of reads in cluster. The pie chart area indicates the relative size of the cluster. The number next to the node denotes the sub-dataset, for which the cluster was obtained. Two clusters are connected with a node if they belong to two different sub-datasets and share at least 25% of their content. The groups of size five (the set of five fully connected pie-charts) represent groups of stable clusters.

Clusters are connected if they were marked as similar and thus shared more then 25% of their content. We looked for sub-graphs, of size five for which all five nodes would be mutually connected. That would mean that all five clusters are coming from the different subsets and share a significant (at least 25% out of 34% possible) number of reads. These groups of clusters (here and after called the stable groups) represent reads that are clustered the same way regardless of the subset of reads selected. Clusters belonging to the stable groups are called the stable clusters. The proportion of reads in the stable clusters was comparable among datasets and equaled on average 64%. As displayed in Fig. 5, we found seven groups of stable clusters. Four groups of stable clusters have clusters with more than 1 000 reads, and two of those four are represented by clusters enriched with *M. oxyfera* or *K. stuttgartiensis/B. sinica* reads. In Table 3 we display the content and the number of reported clusters after the classification procedure for each of the five subsets.

**Table 3.** Subsets information and clustering results.

| Subset | 1 | 2 | 3 | 4 | 5 |
| --- | --- | --- | --- | --- | --- |
| number of *M.oxyfera* reads | 1 499 | 1 563 | 1 528 | 1 544 | 1 529 |
| number of *K.stuttgartiensis/B.sinica* reads | 949 | 918 | 981 | 935 | 906 |
| Clusters after the classification procedure | 14 | 11 | 13 | 13 | 12 |
| Big (>1000 reads) clusters | 5 | 5 | 5 | 5 | 5 |
| % of reads in stable clusters | 65.96 | 64.12 | 61.98 | 64.46 | 64.16 |

Once we estimated the robustness of the classification procedure, we selected the subset that yielded the lowest number of clusters (subset 2, 11 clusters) for downstream analysis. The content of all clusters that were not reported as stable were merged into one cluster. Thus, the original 10 000 reads were spread among 8 clusters. These clusters were used as a classifier for the remaining 21 757 reads in the dataset (Table 4).

**Table 4.** Results of bio reactor metagenome reads classification

| Cluster | Stable | Reads before extension | Reads after extension |
| --- | --- | --- | --- |
| 1 | Yes | 403 | 1 038 |
| 2 | Yes | 168 | 528 |
| 3 | Yes | 1 133 | 3 204 |
| 4 | Yes | 1 540 | 5 151 |
| 5 | Yes | 1 004 | 3 337 |
| 6 | Yes | 181 | 506 |
| 7 | Yes | 1 983 | 6 459 |
| 8 | No | 3 588 | 11 534 |

### 4.5. Assembly of the bio reactor metagenome before and after reads binning

We assembled reads belonging to different clusters separately, and compared the resulting contigs with the results of the assembly of the entire dataset. The total number of contigs after assembly of the partitioned dataset was comparable to the amount of contigs obtained from the assembly of the entire dataset (Table 5). The same can be said about the total length of contigs and contigs length distributions (see supplementary materials). These results, showing that the database partitioning did not lead to the change of the contigs number or their lengths, can be seen as indirect evidence proving that our *k-*mer based binning of metagenome reads results in species-based clustering.

**Table 5.** Results of entire and partitioned bio reactor sequencing data assembly and comparison of obtained contigs to the *M.oxyfera* genome.

| Dataset assembled | Entire dataset | Cluster 1 | Cluster 2 | Cluster 3 | Cluster 4 | Cluster 5 | Cluster 6 | Cluster 7 | Cluster 8 |
| --- | --- | --- | --- | --- | --- | --- | --- | --- | --- |
| Assembly length, bp | 3 251 357 | 5 438 | 10 747 | 380,905 | 377 792 | 601 065 | 0 | 1 602 878 | 41 310 |
| Contigs | 196 | 1 | 1 | 28 | 30 | 47 | 0 | 79 | 4 |
| Contigs mapped on *M.oxyfera* genome | 91 | 0 | 0 | 9 | 1 | 2 | 0 | 71 | 2 |
| Length of mapped contigs, bp | 1 842 182 | 0 | 0 | 132 863 | 11 945 | 21 105 | 0 | 1 497 132 | 17 013 |
| % of *M.oxyfera* genome covered | 54 | 0 | 0 | 1.2 | 0.1 | 0.15 | 0 | 51 | 0.2 |

We compared the assembled contigs obtained for the entire and partitioned datasets to the reference genomes of *M. oxyfera*, *K. stuttgartiensis* and *B. sinica*. Even though we could successfully map around 9% of the reads to the reference genomes of *K. stuttgartiensis* and *B. sinica*, we did not get contigs that could be mapped to these genomes. However, the contigs assembled from the entire and partitioned datasets did map to *M. oxyfera* genome. Only 91 out of 196 contigs obtained from the entire dataset assembly could be mapped back to the *M. oxyfera* genome covering 54% of its length. For the assembly of the partitioned dataset, 85 contigs were mapped to the genome of *M. oxyfera* in total, covering 52.65% of its length. The vast majority of those contigs (79, covering 51% of the *M. oxyfera* genome length) derived from the assembly of reads belonging to one cluster. Thus, our dataset partitioning binned the majority of contigs according to their origin.

## DISCUSSIONS

We described a new approach for efficient, alignment-free binning of metagenomic sequencing reads based on *k*-mer frequencies. Our method successfully classifies reads per organism of origin, for both simulated and real metagenomics data.

As shown in the results section, the approach was used to classify reads obtained by PacBio sequencing of a real bio reactor metagenome. The absolute majority of the reads with known origin (*M. oxyfera* or *K. stuttgartiensis/B. sinica*) were clustered together per origin after pairwise comparison of their *k-*mer profiles and subsequent density-based cluster detection. This result was robust, as we observed during the analysis of five subsets of the original PacBio sequencing data with overlapping content. The same experiment demonstrated that each subset provides a similar number of clusters. Reads with unknown origin tended to cluster similarly among different subsets, again confirming the clustering consistency. Although the majority of reads in the analyzed metagenome was of unknown origin, the results can be used to estimate the microbial community complexity for its most abundant inhabitants.

The binning of the bio-reactor metagenomics dataset had almost no influence on the results of the metagenome assembly. The number of contigs and their lengths obtained for the entire and partitioned datasets were comparable. This indicates that the *k*-mer based reads binning leads to the organism-based partitioning of metagenomic data. Furthermore, contigs, belonging to the same organism, were automatically grouped together when assembling the dataset subjected to the classification procedure. Thus, our *k*-mer based binning technique can be used to interpret metagenomic assembly results. Performing the binning procedure on an artificially generated PacBio datasets lead to a reads classification per organism, even after adding reads with unknown origin (noise reads). Moreover, increasing the proportion of noise reads leads to a better separation between them and the reads with known origin. This observation supports the central hypothesis of this research, namely that *k*-mer distances can be used to cluster reads of the same origin together once those reads provide sufficient coverage of the organisms’ genome.

The main disadvantages of the current implementation of our method is the limited number of reads (10 000) that can be analyzed. As mentioned before, reads, derived from the same organism, will cluster together, but this is possible only under the condition that the organisms’ genome is sufficiently covered. Thus, the described technique is unsuitable for the analysis of metagenomes with a large number of inhabitants or when the inhabitants have large genomes, as 10 000 reads will not be enough to provide sufficient coverage. The depth of the classification that can be performed by the suggested method is still to be discovered.

We believe that adapting our metagenomics reads binning technique for larger sets of data and further investigation of its metagenome resolving capacity would allow to expand the current limits of microbiology in the future.

## CONCLUSIONS

In this study we demonstrated the possibility to detect substructures within a single metagenome operating only with the information derived from the sequencing reads. Results obtained for both artificial and real metagenomic data indicated the reads clustering per their known origin. We have shown the robustness of the obtained results by adding different proportions of “noise” reads to the artificially generated metagenomic data and by comparing the results of binning procedure performed on the different subsets of the same real metagenomic dataset. The obtained results are highly important as they establish a principle that might potentially greatly expand the toolkit for the detection and investigation of previously unknow microorganisms.

## LIST OF ABBREVIATIONS

PacBio: Pacific Biosciences
NGS: next-generation sequencing
N-DAMO: Nitrite/Nitrate dependent Anaerobic Methane Oxidation Annamox - anaerobic ammonium oxidation
WGS: whole-genome shotgun sequencing.

## DECLARATIONS

### Ethics approval and consent to participate

Since in this research no human material or clinical records of patients or volunteers were used, this research is out of scope for a medical ethical committee. This information was verified by the Leiden University Medical Center Medical Ethical Committee.

### Consent for publication

Not applicable

### Availability of data and material

Sequencing reads of bioreactor metagenome were submitted to NCBI under the BioProject number PRJNA487927. Artificial PacBio metagenomic reads with the addition of 0%, 5%, 10%, and 15% of real “noise” reads were submitted to NCBI under the BioProject number PRJNA533970. Supplementary materials (Additional file 1) were deposited on Figshare and available for downloading using the following link: https://doi.org/10.6084/m9.figshare.c.4218857.v1. Example of the classification procedure can be found using the following link: https://git.lumc.nl/l.khachatryan/pacbio-meta/blob/master/analysis/real_data/tsne_subset2/analysis_example.ipynb

### Authors’ contributions

LK algorithm developing, data acquisition, analysis and interpretation, manuscript drafting; SYA conception, data acquisition and analysis, manuscript editing; RHAMV data acquisition, manuscript editing; JFJL conception, manuscript editing, general supervision.

## Acknowledgements

Authors would like to thank the group of Prof. Huub Op den Camp for the bioreactor metagenome material, Prof. Boudewijn P. F. Lelieveldt for the idea to perform dimensional reduction using t-SNE, and Martijn Vermaat for the help with coding.

## ADDITIONAL FILES

Additional file 1: Article supplement (PDF 143 kb).

Section S1: Threshold for the contig-genome similarity using LAST; Section S2: Detailed results of artificial metagenomes binning.

## REFERENCES

[1] Bikel S, Valdez-Lara A, Cornejo-Granados F, Rico K, Canizales-Quinteros S, Soberon X, et al. Combining metagenomics, metatranscriptomics and viromics to explore novel microbial interactions: towards a systems-level understanding of human microbiome. Computational and structural biotechnology journal 2015;13:390–401.

[2] Gosalbes MJ, Abellan JJ, Durban A, Perez-Cobas AE, Latorre A, and Moya A. Metagenomics of human microbiome: beyond 16S rDNA. Clinical Microbiology and Infection 2012;18:47–49.

[3] Maccaferri S, Biagi E, and Brigidi P. Metagenomics: key to human gut microbiota. Digestive diseases 2011;29(6):525–530.

[4] Martin R, Miquel S, Langella P, and Bermudez-Humaran LG. The role of metagenomics in understanding the human microbiome in health and disease. Virulence 2014;5(3):413–423.

[5] Edmonds-Wilson SL, Nurinova NI, Zapka CA, Fierer N, and Wilson M. Review of human hand microbiome research. Journal of dermatological science 2015;80(1):3–12.

[6] Blum HE. The human microbiome. Advances in medical sciences 2017;62(2):414–420.

[7] Holmes E, Li JV, Marchesi JR, and Nicholson JK. Gut microbiota composition and activity in relation to host metabolic phenotype and disease risk. Cell metabolism 2012;16(5):559–564.

[8] Bhatt AP, Redinbo MR, and Bultman SJ. The role of the microbiome in cancer development and therapy. CA: a cancer journal for clinicians 2017;67(4):326–344.

[9] Cho I and Blaser MJ. The human microbiome: at the interface of health and disease. Nature Reviews Genetics 2012;13(4):260.

[10] Sonnenburg JL and Backhed F. Diet–microbiota interactions as moderators of human metabolism. Nature 2016;535(7610):56.

[11] Mullish BH, Marchesi JR, Thursz MR, and Williams HRT. Microbiome manipulation with faecal microbiome transplantation as a therapeutic strategy in clostridium difficile infection. QJM: An International Journal of Medicine 2014;108(5):355–359.

[12] Moloney RD, Desbonnet L, Clarke G, Dinan TG, and Cryan JF. The microbiome: stress, health and disease. Mammalian Genome 2014;25(1-2):49–74.

[13] Contreras AV, Cocom-Chan B, Hernandez-Montes G, Portillo-Bobadilla T, and Resendis-Antonio O. Host-microbiome interaction and cancer: Potential application in precision medicine. Frontiers in physiology 2016;7:606.

[14] He C, Shan Y, and Song W. Targeting gut microbiota as a possible therapy for diabetes. Nutrition Research 2015;35(5):361–367.

[15] Marx CJ. Can you sequence ecology? metagenomics of adaptive diversification. PLoS biology 2013;11(2):e1001487.

[16] Hiraoka S, Yang C-C, and Iwasaki W. Metagenomics and bioinformatics in microbial ecology: current status and beyond. Microbes and environments 2016;31(3):204–212.

[17] Xing M-N, Zhang X-Z, and Huang H. Application of metagenomic techniques in mining enzymes from microbial communities for biofuel synthesis. Biotechnology advances 2012;30(4):920–929.

[18] Tiwari R, Nain L, Labrou NE, and Shukla P. Bioprospecting of functional cellulases from metagenome for second generation biofuel production: a review. Critical reviews in microbiology 2018;44(2):244–257.

[19] Sommer MOA, Church GM, and Dantas G. A functional metagenomic approach for expanding the synthetic biology toolbox for biomass conversion. Molecular systems biology 2010;6(1):360.

[20] Bokulich NA, Lewis ZT, Boundy-Mills K, Mills DA. A new perspective on microbial landscapes within food production. Current opinion in biotechnology 2016;37:182–189.

[21] Trindade M, van Zyl LJ, Navarro-Fernandez J, and Abd Elrazak A. Targeted metagenomics as a tool to tap into marine natural product diversity for the discovery and production of drug candidates. Frontiers in microbiology 2015;6:890.

[22] Zhang MM, Qiao Y, Ang EL, and Zhao H. Using natural products for drug discovery: the impact of the genomics era. Expert opinion on drug discovery 2017;12(5):475–487.

[23] Techtmann SM and Hazen TC. Metagenomic applications in environmental monitoring and bioremediation. Journal of industrial microbiology & biotechnology 2016;43(10):1345–1354.

[24] Kunin V, Copeland A, Lapidus A, Mavromatis K, and Hugenholtz P. A bioinformatician’s guide to metagenomics. Microbiology and molecular biology reviews 2008;72(4):557–578.

[25] Mande SS, Mohammed MH, and Ghosh TS. Classification of metagenomic sequences: methods and challenges. Briefings in bioinformatics 2012;13(6):669–681.

[26] Ding X, Cheng F, Cao C, and Sun X. Dectico: an alignment-free supervised metagenomic classification method based on feature extraction and dynamic selection. BMC Bioinformatics 2015 16(1):323.

[27] Cui H and Zhang X. Alignment-free supervised classification of metagenomes by recursive svm. BMC Genomics 2013;14(1):641.

[28] Liao W, Ren J, Wang K, Wang S, Zeng F, Wang Y, and Sun F. Alignment-free transcriptomic and metatranscriptomic comparison using sequencing signatures with variable length Markov chains. Scientific reports 2016;6:37243.

[29] Laczny CC, Kiefer C, Galata V, Fehlmann T, Backes C, and Keller A. Busybee web: metagenomic data analysis by bootstrapped supervised binning and annotation. Nucleic acids research 2017;45(W1):W171–W179.

[30] Wang Y, Hu H, and Li X. Mbmc: An effective Markov chain approach for binning metagenomic reads from environmental shotgun sequencing projects. Omics: a journal of integrative biology 2016;20(8):470–479.

[31] Kotamarti RM, Hahsler M, Raiford D, McGee M, and Dunham MaH. Analyzing taxonomic classification using extensible Markov models. Bioinformatics 2010;26(18):2235–2241.

[32] Seok H-S, Hong W, and Kim J. Estimating the composition of species in metagenomes by clustering of next generation read sequences. Methods 2014;69(3):213–219.

[33] Lemos LN, Fulthorpe RR, Triplett EW, and Roesch LFW. Rethinking microbial diversity analysis in the high throughput sequencing era. Journal of microbiological methods 2011;86(1):42– 51.

[34] Janssen P, Goldovsky L, Kunin V, Darzentas N, and Ouzounis CA. Genome coverage, literally speaking: The challenge of annotating 200 genomes with 4 million publications. EMBO reports 2005; 6(5):397–399.

[35] Akondi KB and Lakshmi VV. Emerging trends in genomic approaches for microbial bioprospecting. Omics: a journal of integrative biology 201317(2):61–70.

[36] Hunter-Cevera JC. The value of microbial diversity. Current Opinion in Microbiology 1998;1(3):278–285.

[37] Pace NR. Mapping the tree of life: progress and prospects. Microbiology and molecular biology reviews 2009;73(4):565–576.

[38] Grattepanche J-D, Santoferrara LF, McManus GB, and Katz LA. Diversity of diversity: conceptual and methodological differences in biodiversity estimates of eukaryotic microbes as compared to bacteria. Trends in microbiology 2014;22(8):432–437.

[39] Zinger L, Gobet A, and Pommier T. Two decades of describing the unseen majority of aquatic microbial diversity. Molecular Ecology 2012;21(8):1878–1896.

[40] Szalkai B, Scheer I, Nagy K, Vertessy BG, Grolmusz V. The metagenomic telescope. PloS One 2014;9(7):e101605.

[41] Rosen GL, Polikar R, Caseiro DA, Essinger SD, and Sokhansanj BA. Discovering the unknown: improving detection of novel species and genera from short reads. BioMed Research International 2011;2011:495849. doi: 10.1155/2011/495849.

[42] Kislyuk A, Bhatnagar S, Dushoff J, and Weitz JS. Unsupervised statistical clustering of environmental shotgun sequences. BMC Bioinformatics 2009;10(1):316.

[43] Roumpeka DD, Wallace RJ, Escalettes F, Fotheringham I, and Watson M. A review of bioinformatics tools for bio-prospecting from metagenomic sequence data. Frontiers in genetics 2017;8:23.

[44] Wu Y-W and Ye Y. A novel abundance-based algorithm for binning metagenomic sequences using l-tuples. Journal of Computational Biology 2011;18(3):523–534.

[45] Wang Y, Leung HCM, Yiu S-M, and Chin FYL. Metacluster 4.0: a novel binning algorithm for NGS reads and huge number of species. Journal of Computational Biology 2012;19(2):241–249.

[46] Van Lang T, Van Hoai T, et al. A two-phase binning algorithm using l-mer frequency on groups of non-overlapping reads. Algorithms for Molecular Biology 2015;10(1):2.

[47] Song K, Ren J, Reinert G, Deng M, Waterman MS, and Sun F. New developments of alignment-free sequence comparison: measures, statistics and next-generation sequencing. Briefings in bioinformatics 2013;15(3):343–353.

[48] Girotto S, Pizzi C, and Comin M. Metaprob: accurate metagenomic reads binning based on probabilistic sequence signatures. Bioinformatics 2016;32(17):i567–i575.

[49] Anvar SY, Khachatryan L, Vermaat M, van Galen M, Pulyakhina I, Ariyurek Y, Kraaijeveld K, den Dunnen JT, de Knijff P, Ac’t Hoen P, et al. Determining the quality and complexity of next-generation sequencing data without a reference genome. Genome Biology 2014;15(12):555.

[50] Ono Y, Asai K, and Hamada M. Pbsim: PacBio reads simulator—toward accurate genome assembly. Bioinformatics 2012;29(1):119–121.

[51] van Eijk E, Anvar SY, Browne HP, Leung WY, Frank J, Schmitz AM, Roberts AP, and Smits WK. Complete genome sequence of the Clostridium difficile laboratory strain 630d erm reveals differences from strain 630, including translocation of the mobile element ctn 5. BMC Genomics 2015;16(1):31.

[52] Haroon MF, Hu S, Shi Y, Imelfort M, Keller J, Hugenholtz P, Yuan Z, and Tyson GW. Anaerobic oxidation of methane coupled to nitrate reduction in a novel archaeal lineage. Nature 2013;500(7464):567.

[53] Chaisson MJ and Tesler G. Mapping single molecule sequencing reads using basic local alignment with successive refinement (blasr): application and theory. BMC Bioinformatics 2012;13(1):238.

[54] Altschul SF, Gish W, Miller W, Myers EW, and Lipman DJ. Basic local alignment search tool. Journal of molecular biology 1990;215(3):403–410.

[55] Chin C-S, Peluso P, Sedlazeck FJ, Nattestad M, Concepcion GT, Clum A, Dunn C, O’Malley, Figueroa-Balderas RR, Morales-Cruz A, et al. Phased diploid genome assembly with single-molecule real-time sequencing. Nature methods 2016;13(12):1050.

[56] Hamada M, Ono Y, Asai K, and Frith MC. Training alignment parameters for arbitrary sequencers with last-train. Bioinformatics 2016;33(6):926–928.

[57] van der Maaten L and Hinton G. Visualizing data using t-sne. Journal of machine learning research 2008;9(Nov):2579–2605.

[58] Ester M, Kriegel H-P, Sander J, Xu X, et al. A density-based algorithm for discovering clusters in large spatial databases with noise. In Kdd 1996;96:226–231.

[59] Frank J, Lucker S, Vossen RHAM, Jetten MSM, Hall RJ, Op den Camp HJM, and Anvar SY. Resolving the complete genome of Kuenenia Stuttgartiensis from a membrane bioreactor enrichment using single-molecule real-time sequencing. Scientific reports 2018;8(1):4580.

