## Additional file 1 for "Reference-free resolution of long-read metagenomic data"

*Supplementary Materials*

**Section S1. Threshold for the contig-genome similarity using LAST**

Since all of the aligned contigs were expected to have some similarity to the reference genome, it was necessary to determine the similarity cut-off separating irrelevant results. To determine this threshold, the curve representing the number of contigs versus the similarity to the reference genome was analyzed.

We assumed that the number of irrelevant contigs will drop with the increasing of the similarity cut-off faster than the number of contigs that have true affinity to the genome. Indeed, we noticed, that the left part of the curve was decreasing faster in comparison with the right part (See Figure S1). Thus, the inflection point at 12%, dividing the fast declining part of the curve from the slowly declining part, was chosen as a threshold.

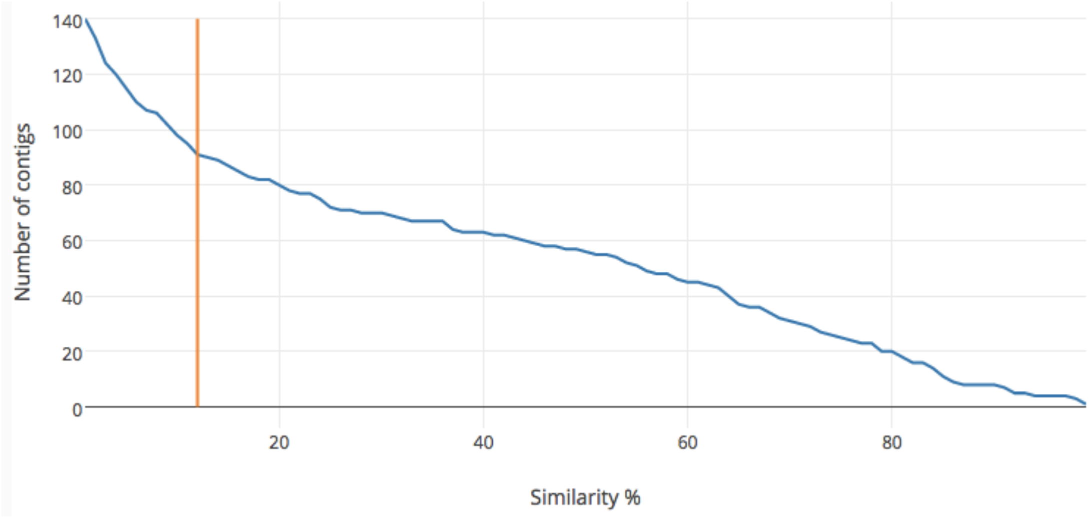


Figure S1. The dependency of the number of contigs mapped on *M. oxyfera* genome from the similarity %. The chosen similarity cut-off is shown as a vertical red line.

**Section S2. Detailed results of artificial metagenomes binning.**

Results for 0% noise, 5% noise, 10% noise and 15% noise datasets are presented in the Tables S1, S2, S3 and S4 correspondingly.

| Cluster | Number of reads | | | | | |
|---|---|---|---|---|---|---|
| | *S. mitis* | *P. acnes* | *S. epidermidis* | *A. calcoaceticus* | *P.aeruginosa* | noise |
| 1 | 0 | 0 | 0 | 0 | 3086 | - |
| 2 | 1214 | 0 | 41 | 14 | 0 | - |
| 3 | 0 | 1443 | 0 | 5 | 185 | - |
| 4 | 0 | 0 | 0 | 0 | 356 | - |
| 5 | 26 | 0 | 1385 | 25 | 0 | - |
| 6 | 6 | 0 | 22 | 2192 | 0 | - |

Table S1. Content of clusters after binning 0% noise dataset

| Cluster | Number of reads | | | | | |
|---|---|---|---|---|---|---|
| | *S. mitis* | *P. acnes* | *S. epidermidis* | *A. calcoaceticus* | *P.aeruginosa* | noise |
| 1 | 0 | 5 | 0 | 0 | 3222 | 30 |
| 2 | 38 | 1366 | 37 | 22 | 222 | 459 |
| 3 | 1119 | 0 | 0 | 0 | 0 | 0 |
| 4 | 2 | 0 | 24 | 2089 | 2 | 3 |
| 5 | 24 | 0 | 1315 | 14 | 0 | 7 |

Table S2. Content of clusters after binning 5% noise dataset

| Cluster | Number of reads | | | | | |
|---|---|---|---|---|---|---|
| | *S. mitis* | *P. acnes* | *S. epidermidis* | *A. calcoaceticus* | *P.aeruginosa* | noise |
| 1 | 0 | 1009 | 0 | 0 | 11 | 4 |
| 2 | 0 | 286 | 0 | 0 | 162 | 399 |
| 3 | 0 | 0 | 0 | 0 | 300 | 0 |
| 4 | 0 | 0 | 0 | 0 | 2783 | 19 |
| 5 | 18 | 0 | 1251 | 10 | 0 | 11 |
| 6 | 2 | 0 | 14 | 1981 | 0 | 8 |
| 7 | 1091 | 0 | 11 | 11 | 0 | 0 |
| 8 | 10 | 3 | 28 | 11 | 8 | 559 |

Table S3. Content of clusters after binning 10% noise dataset

| Cluster | Number of reads | | | | | |
|---|---|---|---|---|---|---|
| | *S. mitis* | *P. acnes* | *S. epidermidis* | *A. calcoaceticus* | *P.aeruginosa* | noise |
| 1 | 0 | 0 | 0 | 0 | 2875 | 7 |
| 2 | 985 | 0 | 22 | 0 | 0 | 0 |
| 3 | 0 | 763 | 0 | 0 | 4 | 14 |
| 4 | 17 | 0 | 1181 | 4 | 0 | 18 |
| 5 | 20 | 1 | 5 | 1868 | 57 | 7 |
| 6 | 0 | 44 | 0 | 0 | 70 | 675 |
| 7 | 4 | 0 | 8 | 9 | 0 | 762 |

Table S4. Content of clusters after binning 15% noise dataset
